# Assessment of the adverse effects of pollution on farmland bird diversity in contemporary agricultural landscapes

**DOI:** 10.1101/2025.03.23.644816

**Authors:** Maitreyi Sur, David Kleijn, Merel Soons, Ruud Foppen, Caspar A. Hallmann, Eelke Jongejans, Leo Posthuma, Henk Sierdsema, Jaap Slootweg, Chris van Turnhout, Hans de Kroon

## Abstract

The escalating global demand for food has intensified agricultural practices, leading to substantial changes in land use. This transformation poses a threat to farmland biodiversity, compounded by the presence of pollutants from anthropogenic activities. While the impact of specific pollutants is known in controlled environments, their compounded effects under field conditions remain largely unexplored. We investigated correlations between farmland bird distribution, landscape features, land use patterns, and anthropogenic pressures, including nutrient pollution, acidifying compounds, and synthetic chemicals. Using distribution maps of the Netherlands at a 1×1 km² grid cell, we analyzed the association of farmland bird species richness and abundance with landscape characteristics and varying levels of exposures to unintended pollutants. We also compared species richness distribution patterns between 1998 and 2018. We found a strong negative relationship between farmland bird species richness and abundance with atmospheric deposition of inorganic nitrogen (NHx, NOy). Furthermore, mixed associations were observed between farmland birds and local toxic pressure variation in surface waters, with consistent relationships to industrial chemicals (negative) and products of combustion (positive). Lastly, change in species richness from 1998 and 2018, showed that many of the relationships observed now were already evident two decades ago, with recent declines in species richness concentrated in landscapes hosting a considerable number of species, and low nitrogen deposition grid cells. We conclude that although it is likely that there is some direct negative effects of pollutants on farmland birds, it is reasonable to also assume that the identified relationships are proxies for the overarching intensity of farming, human disturbance, and broader landscape changes. Our study highlights a) a possible role of synthetic pollutants and acidifying eutrophicating substances in farmland bird decline b) the need for well-designed field studies to complement correlative evidence from big data approaches such as ours to enhance our understanding and c) the broader implications for sustainable land management, emphasizing the importance of a holistic approach in addressing the intricate relationships between pollutants and landscape changes.

## 1 Introduction

Growing food demand has driven the transformation of agricultural landscapes worldwide (IPBES, 2019). To meet the needs of a rapidly expanding global population and fueled by global trade, agricultural practices have intensified, leading to changes in land use, increased mechanization, and the expansion of monoculture farming (Mauser et al., 2015; Tilman et al., 2011). The cumulative effects of intensive farming on farmland biodiversity are concerning, as they contribute to the ongoing decline of many species and the homogenization of landscapes (Campbell et al., 2017; Emmerson et al., 2016; Newbold et al., 2015). The effects on biodiversity are exacerbated by the presence of pollutants resulting from agricultural and other anthropogenic activities (Czyżewski et al., 2020). These pollutants, including pesticides, fertilizers, organic compounds, and other chemical compounds, can act as stressors, disrupting ecological processes and exerting adverse effects on various organisms within farmland ecosystems (Köhler & Triebskorn, 2013; Stevens et al., 2020). Even though the mode of action of specific pollutants to target species may be known through laboratory experimentations, the compounding effects under field conditions remain largely unexplored, mainly due to the lack of comprehensive geographic monitoring (Köhler & Triebskorn, 2013; Tang et al., 2021) and practical problems encountered when evaluating unintended mixture exposures.

Presently, there is a heated debate on the relationship between pollutants and biodiversity and its implications for society. For instance, in the Netherlands, economic activities that increase emission of nitrogen compounds, such as from farming or construction, are currently prohibited because of the deleterious impact of excessive nitrogen (N) deposition on species and habitats that are protected under (inter)national laws, including EU’s Bird and Habitat Directives. In response, the sectors and actors that are most strongly affected are publicly challenging the necessity of any restrictions. The evidence base on the subject is rapidly expanding. However, studies that examine the relationships between biodiversity and various stressors simultaneously at large spatial scales remain relatively scarce.

To help fill this knowledge gap, in this study we link country-wide, high-resolution farmland bird population distribution data to an extensive, high-quality spatial dataset of pollutant levels in the Netherlands. Research on avian populations, in particular farmland birds, has seen a notable increase in recent times as a means of studying the impacts of pollutants such as pesticides and excessive nutrients in agricultural landscapes (Donald et al., 2001; Hallmann et al., 2014; Moreau et al., 2022; Newton, 2004). Composite indicators such as species richness and abundance of farmland birds could potentially reflect population responses, but also mirror effects of ecosystem health on a species assemblage (Gregory & van Strien, 2010). Studies on individual species such as the Corn Bunting (*Emberiza calandra*; Brickle & Harper, 2002; Brickle et al., 2000), Yellowhammer (*Emberiza citrinella*; Morris et al., 2005), Patridge (*Perdix perdix*; Aebischer & Ewald, 2004) and Skylark (*Alauda arvensis*; Engel et al., 2012) have provided a strong foundation for investigating the effects of pollutants on groups of species with similar functional traits (e.g. similar food and habitat preferences).

Most studies at the local as well as continental level quantify pollutants using economic indicators collected through sales monitoring or farming census (e.g., amount purchased, amount used; Li et al., 2020; Rigal et al., 2023). While this provides important insights, it is questionable whether this is an accurate indicator of the levels of pollutants that birds are actually exposed at the local scale. The complexity and heterogeneity of present-day agricultural landscapes, and their proximity to urban areas, probably results in considerable spillover of pollutants to and from different parts of the agricultural landscape. For example, pollutants in surface water resulting from agricultural runoff, industrial discharges, and other anthropogenic activities can diffuse from point sources (as recorded from farm census data), to lower regions of a watershed. Similarly, nearly all pollutants emitted into the atmosphere from industrial emissions, agricultural activities, and combustion processes end up as dry and wet depositions on land surfaces at varying distances away from the source of origin.

Pollutant diffusion in watersheds may expose avian species in different parts of a landscape, and may lead to reduced reproductive success, impaired immune function, and altered behavior in farmland birds (Humann-Guilleminot et al., 2019). Several studies have also shown adverse effects of sulfur and nitrogen deposition on nutrient cycling, soil conditions, vegetation composition and structure and invertebrate communities (Bobbink, 1991; Duprè et al., 2010; Maskell et al., 2010; Stevens et al., 2020; Sutton et al., 2011; Van den Berg et al., 2011). All these changes on agricultural landscapes may result in lasting effects on farmland birds by changing the availability and quality of food resources, nesting possibilities and breeding habitats (Barton et al., 2023; Dise, 2011; Lennon et al., 2019; Nijssen et al., 2017).

The Netherlands is amongst the most intensively farmed countries in the world (OECD, 2023). At the same time, the country faces a high biodiversity loss, particularly for bird species living in agricultural landscapes (‘farmland birds’; Environmental Data Compendium, 2023; Sovon 2018). In the past decades, both agricultural intensification and environmental impacts including species distribution and abundances have been monitored in the Netherlands at high spatial resolution. The Netherlands is on par with the UK, the leading country in comprehensive bird monitoring. In recent years, the Netherlands has furthermore meticulously collected and disseminated geographical and spatial data related to land use and associated environmental indicators (more details on https://www.pdok.nl/) together with detailed satellite imagery and high-resolution aerial photographs (more details on https://www.satellietdataportaal.nl/).The extensive coverage (nationwide) and high quality (complying with national and international standards, including the European INSPIRE standard) and resolution (eg land use data available at 5m resolution) of data accessible in the Netherlands make it uniquely capable of conducting a nationwide analysis of the probable causes of exposure to- and effects of pollution on farmland biodiversity at a national scale.

In this study we ask whether and how the distribution of farmland birds across the country correlates with landscape features, land use patterns, and anthropogenic pressures, including nutrient pollution, acidifying compounds, and diverse synthetic chemicals. We hypothesize that direct and indirect effects of diverse levels of pollutants have differently compromised the vitality of farmland bird populations, leading to lower farmland bird abundances and fewer species in the landscape at higher exposures. We also hypothesize that ground-nesting and non-ground-nesting farmland birds have different relationships with environmental variables and pollutants due to their distinct habitat preferences (open versus closed landscapes) and associated exposure levels. We expect that many ground-nesting farmland birds, often residing in waterbodies (e.g., all waders), are more directly exposed to pollutants in surface water than non-ground-nesting farmland birds. Conversely, non-ground-nesting birds, which are typically found in closed landscapes, are exposed to different levels and types of pollutants due to the farming practices associated with these environments, such as the higher use of insecticides in arable systems compared to grassland-based systems. To test these hypotheses, we first created distribution maps of the Netherlands based on a 1×1 km2 resolution grid cells containing information on farmland bird species richness and abundance, landscape characteristics, airborne pollution estimates and water pollution estimates. For two distinct groups of farmland birds (ground nesting and non-ground nesting), we then analyzed to what extent species richness and abundances were associated with landscape characteristics (such as openness, groundwater level, and proportion of grassland, natural and urban area) and pressures exerted by varying levels of exposures to unintended pollutants. Finally, to examine whether these relationships have changed over time, we compared species richness distribution patterns between 1998 and 2018.

## 2 Methods

### 2.1 Data collection

#### 2.1.1 Bird distribution data

To establish bird distributions and abundances, we used distribution data of 28 farmland bird species in the Netherlands, collected as part of breeding bird distribution atlases Sovon Dutch Centre for Field Ornithology (Sovon, 2018). We made a distinction between species breeding on the ground in open landscapes (n=13 species; hereafter “ground nesters”, including typical meadow birds), and those nesting in shrubs, hedges, trees or buildings in more heterogeneous landscapes (n=15 species; hereafter “non-ground nesters”) (Table 1). Field work for the Atlas was conducted between 2013 to 2015, within standardized 5×5 km atlas grids together covering the entire country. During the breeding season (April - June), bird species presence was scored (i.e., seeing and/or hearing; more details can be found in Vorisek et al.,2008 and Bibby et al., 2012). Migratory species were monitored and handled in similar ways as non-migratory species. Additionally, abundance count scores were collected during two standardized timed visits in a systematic sample of 1×1 km squares. In most cases 8 out of 25 square km grids per 5×5 km atlas square were selected in a systematic manner which resulted in a total of 11680 square km grids(SI1; hereafter referred to as “km grid”; Altenburg et al., 2017), spread over a six-week period, within the breeding season. The visits included either a 5-minute point count or 2 x 5 minute point count that was done from the center of each 1×1 km square, during which all individuals seen were counted. All observations were associated with a column that indicated if the observation came from a 5-minute point count or the first 5-minutes of a 10-minute point count (from here on called “survey-type”). We used this 5-minute point count data to calculate overall and grouped species richness and abundance within all 1×1 km squares.

**Table 1:** List of farmland bird species in the Netherlands. These are species that are dependent on farmlands for feeding and nesting and are not able to thrive in other habitats. We grouped all species into two groups: ground nesting and non-ground nesting birds. Details of these groups are provided in the text.

| Ground nesting Birds | Non-ground nesting birds |
| --- | --- |
| Black-tailed Godwit | Barn Swallow |
| Common Quail | Common Linnet |
| Common Redshank | Common Whitethroat |
| Eurasian Curlew | Common Wood Pigeon |
| Eurasian Oystercatcher | Eurasian Tree Sparrow |
| Eurasian Skylark | European Stonechat |
| Grey Partridge | European Turtle Dove |
| Meadow Pipit | House Martin |
| Northern Lapwing | Icterine Warbler |
| Northern Shoveler | Mistle Thrush |
| Yellow Wagtail | Rook |
|  | Sand Martin |
|  | White Stork |
|  | Yellowhammer |

Additionally, we calculated the change in presence of individual farmland bird species and bird groups in the last 20 years within each 1×1 km square. These calculations were based on an additional data set acquired from a previous breeding bird distribution atlas capturing data during Breeding Bird Atlas (Sovon 2002), for which fieldwork was carried out in 1998-2000 using the same field work protocol (Sovon Vogelonderzoek Nederland, 2002) and conducted in exactly the same systematic sample of 1×1 km squares.

#### 2.1.2 Domains

Soil type, land use and landscape composition may strongly drive variation in bird distribution and abundance and are often spatially autocorrelated, resulting in geographically distinct landscape types. To correct for the confounding effects of such geographical heterogeneity on bird monitoring data across different regions of the Netherlands, we conducted our analyses within areas that can be designated as distinct agricultural landscapes, hereafter referred to as “domains”. The maps of the domains were made by combining different geographical details such as soil types, historical reclamation details, physical geographic properties, etc in a Geographical Information System (GIS) resulting in 21 distinct domains (more details can be found in Kwak & Louwe Kooijmans, 2021 and SI2; also depicted in SI3)

#### 2.1.3 Landcover data

We used the national land use data of the Netherlands for the year 2018 (LGN2018; Hazeu et al. 2018) at a resolution of 5×5 meters, with details of 48 land use sub-classes. We consolidated these into 8 main s use classes namely: arable crops, agricultural grasslands, natural grasslands, heather and moorland, swamps and dunes, forest areas, water areas, and built-up areas (details of each of the classes in provided in SI4). We then calculated the proportion of each land use class within every 1×1 km grid cell.

#### 2.1.4 Openness

We used the openness of a landscape as a variable that describes the landscape structure (more details in SI1). The presence of farmland birds, especially meadow birds, is strongly related to the openness of a landscape. We used openness maps produced for the years 2015 to 2018 by WEnR with the model ViewScape (Meeuwsen and Jochem, 2015). We used GIS to calculate mean openness for these years within each km square grid described above (more details in SI2; depicted in SI5).

#### 2.1.5 Average Spring Groundwater Level (GVG)

Soil moisture and groundwater depth are critical factors that influence the population densities of farmland bird community, especially meadow birds (Kleijn & van Zuijlen, 2004). We used the spatial map of Average Spring Groundwater Level or GVG (more details in SI2) at a scale of 1:250,000 for the year 2018 to calculate average values within a square km grid (depicted in SI5).

#### 2.1.6 Toxic pressure of unintended mixtures (surface water toxicity)

We aimed to collect information on site-specific (1×1 km) data on exposure to synthetic chemicals, which causes variation in toxic pressures. Lacking soil monitoring data for chemical pollutants at the required scale and hypothesizing that concentrations of chemical pollutants in local aquatic systems reflect the regional use of them, we collected aquatic exposure concentration data as a proxy to quantify the toxic pressure of unintended mixtures of chemicals. The toxic pressure metrics were derived for data from the period 2013-2018, recognizing that the same substances were not analyzed at all locations at all times. The toxic pressure metrics were thus calculated per studied compound, as well as for specific subgroups of compounds that are land-use related aggregated metrics for selected compound groups. These were msPAF NHx (Nitrogen emissions from agricultural activities), msPAF pesticides (pesticides related to agriculture), msPAF Industrial (industrial compounds emitted from industrial chemicals), and msPAF combustion (human activities causing combustion). The toxic pressure of the pollutants, is expressed as the toxic pressure of that chemicals, quantified as the Potentially Affected Fraction (hereafter PAF) per chemical, or as multi-substance msPAF (for unintended mixtures). The latter is calculated from the former by aggregation using the concentration addition approach (more details in SI2).

The density of sampling points for chemical analyses was lower than the density needed to match the distribution of the bird data. We therefore used Empirical Bayesian kriging (EBK) to geostatistically interpolate data collected at point locations within the Netherlands (depicted in Fig. 1A). We then used the interpolated maps to calculate the mean local toxic pressure proxies for each of the compound groups within the 1×1 km grids described above.

**Figure 1:**
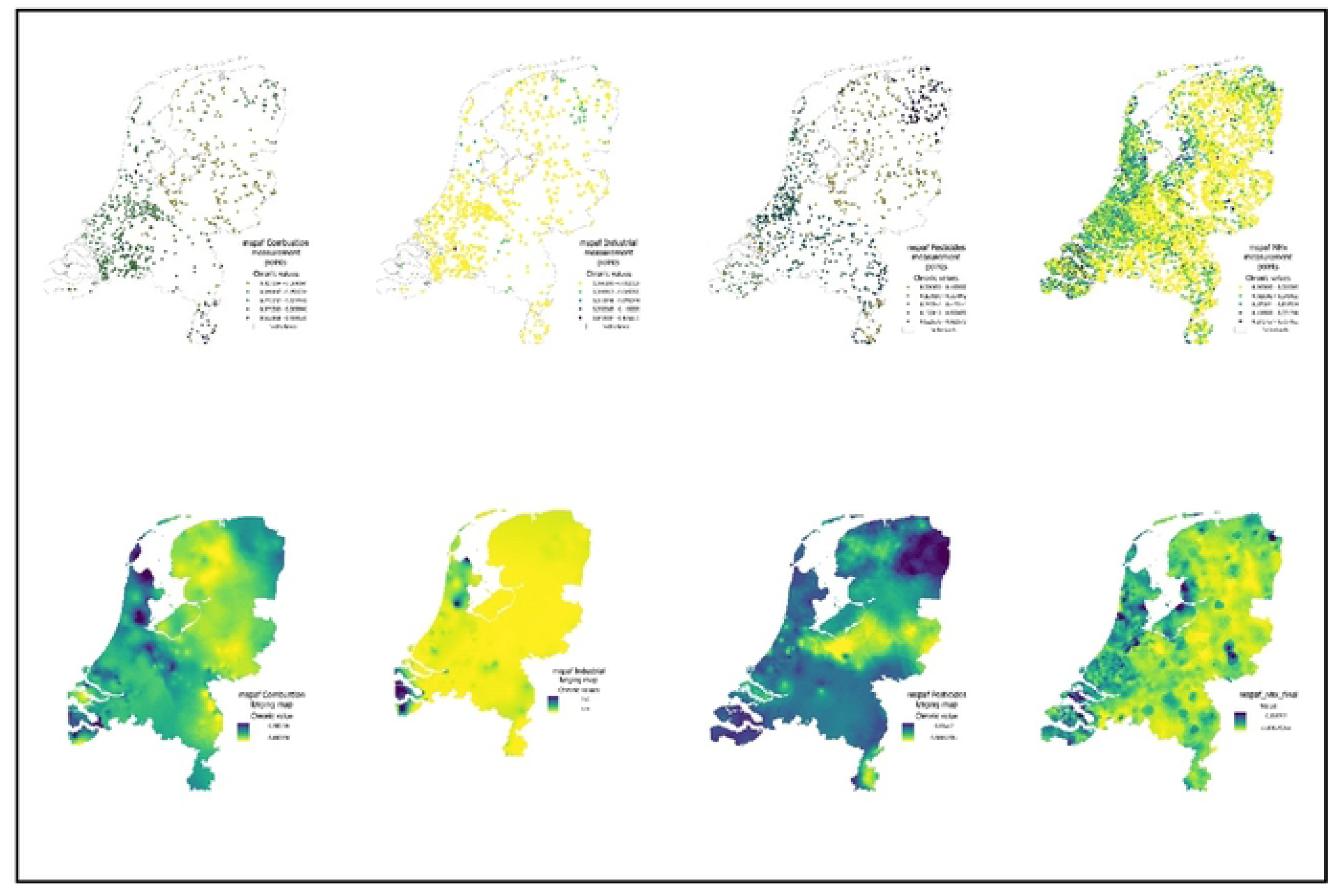
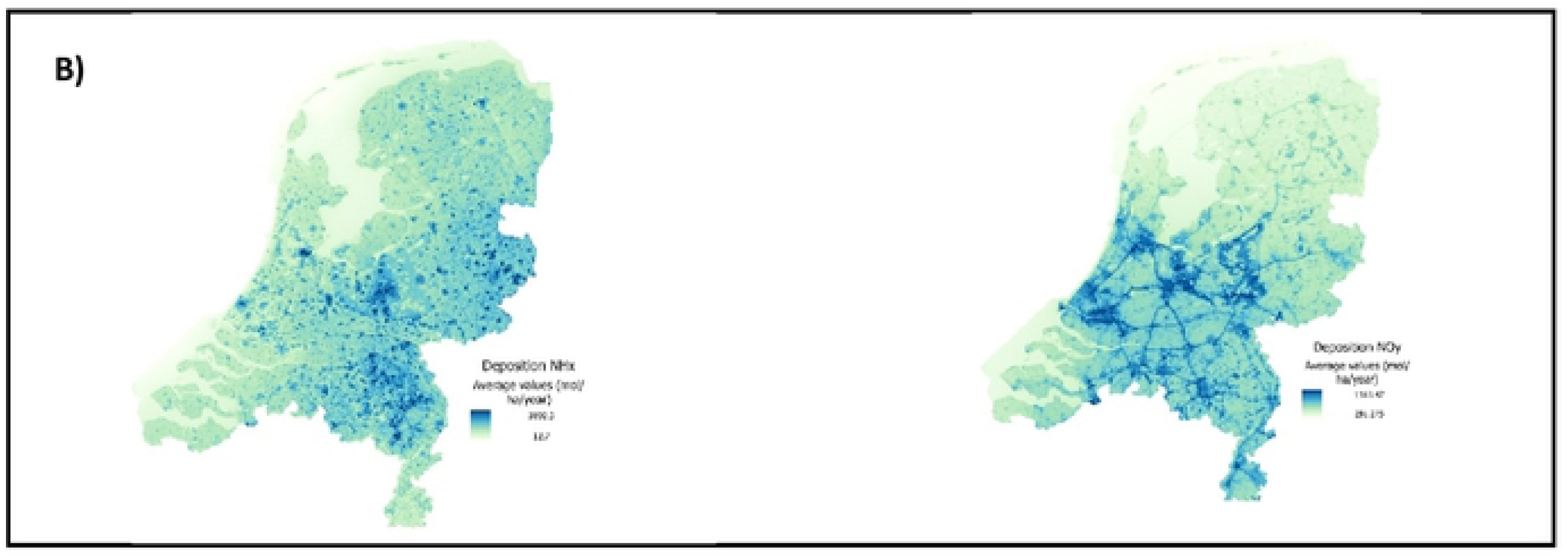
A) Details of toxicity of ammonia/ammonium (NHx), microorganic compounds (Organic compounds) and crop protection compounds (Pesticides) in the surface waters of Netherlands. The top row shows distribution of points where measurements were taken and toxic pressures were calculated for each category of substances between the years 2013-2018. Bottom row shows interpolated surfaces of toxic pressure of each category of substance developed using Empirical Bayesian kriging in ArcGIS pro (details in the text). B) Average atmospheric deposition of NHx and NOy within all 1X1km square grids in the Netherlands between years 2013-2015: Dark colors indicate higher deposition values (in mol/ha/year) while lighter colors indicate lower deposition values. Details included in the main text.

#### 2.1.7 Deposition data of acidifying compounds and nitrogen

The total atmospheric deposition of acidifying contaminants and nitrogen consists of the sum of dry and wet deposition (van der Swaluw et al. 2011). Each year, deposition maps for these compounds are published in the Netherlands on a 1×1 km scale. Maps showing the spatial

distribution of the total annual acidifying and nitrogen deposition in the Netherlands are made by using the combination of measurements and model output from the Operational Priority Substances model (OPS-model), which is described as a variable spatial resolution atmospheric transport and deposition model (van Jaarsveld, 2004). We downloaded deposition maps on acidifying and eutrophicating substances (NOy, NHx) for the years 2013 to 2015. We then calculated the average deposition of NOy and NHx from 2013 to 2015 in all 1×1 km square grids where data on birds were available (depicted in Fig. 1B).

We used ArcGIS pro (version 3.03) for all spatial analysis and calculations.

### 2.2 Analysis

#### 2.2.1 Selecting covariates for inclusion in statistical models

We considered 16 explanatory variables in our linear mixed models. These included the environmental stressors (n=6; msPAF NHx, msPAF pesticides, msPAF industrial, msPAF combustion, deposition of NOy, deposition of NHx) and landscape variables (n=10; proportion of the 1×1 cell covered by arable crops, agricultural grass, natural grasslands, forests, water, shrubs, swamps and dunes, built area, together with openness and average spring groundwater level). We checked for multicollinearity among variables using the variation inflation factor (VIF), with a cut-off value of 3, using models with the structure:

Abundance^1,2^ ∼ All explanatory variables
Species Richness^1,2^ ∼ All explanatory variables
^1,2^ ground nesting and non-ground nesting bird groups

#### 2.2.2 Effects of deposited and toxic chemicals on farmland birds

Focusing on spatial variability, we constructed four separate linear mixed-effects models (package nlme; Pinheiro et al., 2013, R Core Team, 2013) to evaluate how abundance and species richness (response variables) of two separate groups of farmland birds (ground nesting and non-ground nesting) were related to farmland stressors and landscape variables (explanatory variables). We included domains as a random effect in the model structure. When abundance was used as a response variable, we also included species as a random variable in the models to account for differences between species in overall abundance. We also included survey type as a random effect, to account for difference in observation effort. We scaled all explanatory variables within each domain (Gelman 2008).

We then evaluated, for each group of farmland birds, all possible sub-models using the dredge function of the MuMIN package in R (v1.47.1; Doherty et al. 2012; Barton 2019). Sub-models were then ranked based on the Akaike information criterion (AIC) to identify the best supported model explaining the response variable (Anderson & Burnham 2002, Anderson 2007). When no single model had > 90% of model weights, the top supported models (delta AIC<2) were averaged to get model estimates (Anderson & Burnham 2002).

We additionally ran individual models for the abundance of each species separately with all explanatory variables described above and domains and survey type as random effects.

#### 2.2.3 Effects of environmental stressors on changes in species richness from 1998 to 2018

Focusing on temporal trends, to evaluate how changes in species richness from 1998 to 2018 (hereafter ‘species richness trend’) were related to pollutants and landscape variables (explanatory variables described above), we constructed linear mixed-effects models with species richness trend (absolute change in species richness per km sq) for ground nesting and non-ground nesting birds (response variables). It’s important to mention that we did not compute the changes in landscape variables or level of pollutants between 1998 and 2018. Instead, we use the same calculated explanatory variables (landscape variables and pollutants) as calculated in the previous analysis, spanning between the years 2013 to 2018. We again used model selection and model averaging to get model estimates.

#### 2.2.4 Sensitivity tests

To examine how the results were affected by geostatistical interpolations (kriging of msPAF metrics from observation sites to the 1*×*1 grids) we did a separate analysis with the subset of the square kilometer grids, restricting it to grids located within a 5-kilometer radius of measurement locations for water toxicity.

## 3 Results

### 3.1 Characterization of collected data

Farmland bird presence and numbers were available for 11,680 square km grid cells. We omitted grid cells that did not fall within the previously defined domains, which left 8,311 square km grid cells that were used in our analyses. The number of 1*×*1 km^2^ grid cells where farmland birds were monitored varied between the 21 agricultural domains with an average of 424 grids per domain (median = 375, min. = 61, max. = 1125). Species richness and abundance of the two groups of birds showed distinct distribution patterns across various agricultural landscapes. Farmland birds of both groups were present in high abundance and were most diverse in low peat and sea clay polder domains. As expected, bird species that do not nest on the ground showed higher richness within more heterogeneous landscapes that featured a variety of vegetation, including shrubs, bushes, and trees. We included all 16 explanatory variables in our analysis noting that VIF values remained below 3 for all models (SI6). We acknowledge that while some of the pollutants examined in our analysis showed correlations (e.g., mixture toxic pressure attributable to NHx and Pesticides; SI7), it was crucial to include these variables to offer a comprehensive understanding of stressors in agricultural landscapes.

### 3.2 Spatial relations between land use characteristics and pollutants and farmland birds

#### 3.2.1 Ground nesting bird species

Our results showed the expected relations between the abundance and species richness of ground-nesting birds and land use variables. For example, openness of the landscape, the proportion of natural grasslands and groundwater level were positively related with species abundance and richness, while the proportion forest cover was negatively related to abundance and richness of this farmland bird group that contains species like Lapwing, Black-tailed godwit and Skylark (Fig.2a; SI8).

After accounting for the relationships with the land use variables, we found strong support for negative relationships between deposition of NOy and NHx and abundance and species richness of ground nesting farmland birds. Low deposition square km cells (<400 NOy mol ha^-1^ yr^-1^ or <500 NHx mol ha^-1^ yr^-1^, Fig.3,4) harbored between 2 and 4 farmland bird species on average across domains. At the other extreme, in high deposition square km cells (>700 NOy mol ha^-1^ yr^-1^ or >2000 NHx mol ha^-1^ yr^-1^, Fig.4), farmland birds were virtually absent. Individual species models showed negative relationships with deposition of NHx for eight out of 11 species and with NOy deposition for all 11 species (Fig. 3).

**Figure 2:**
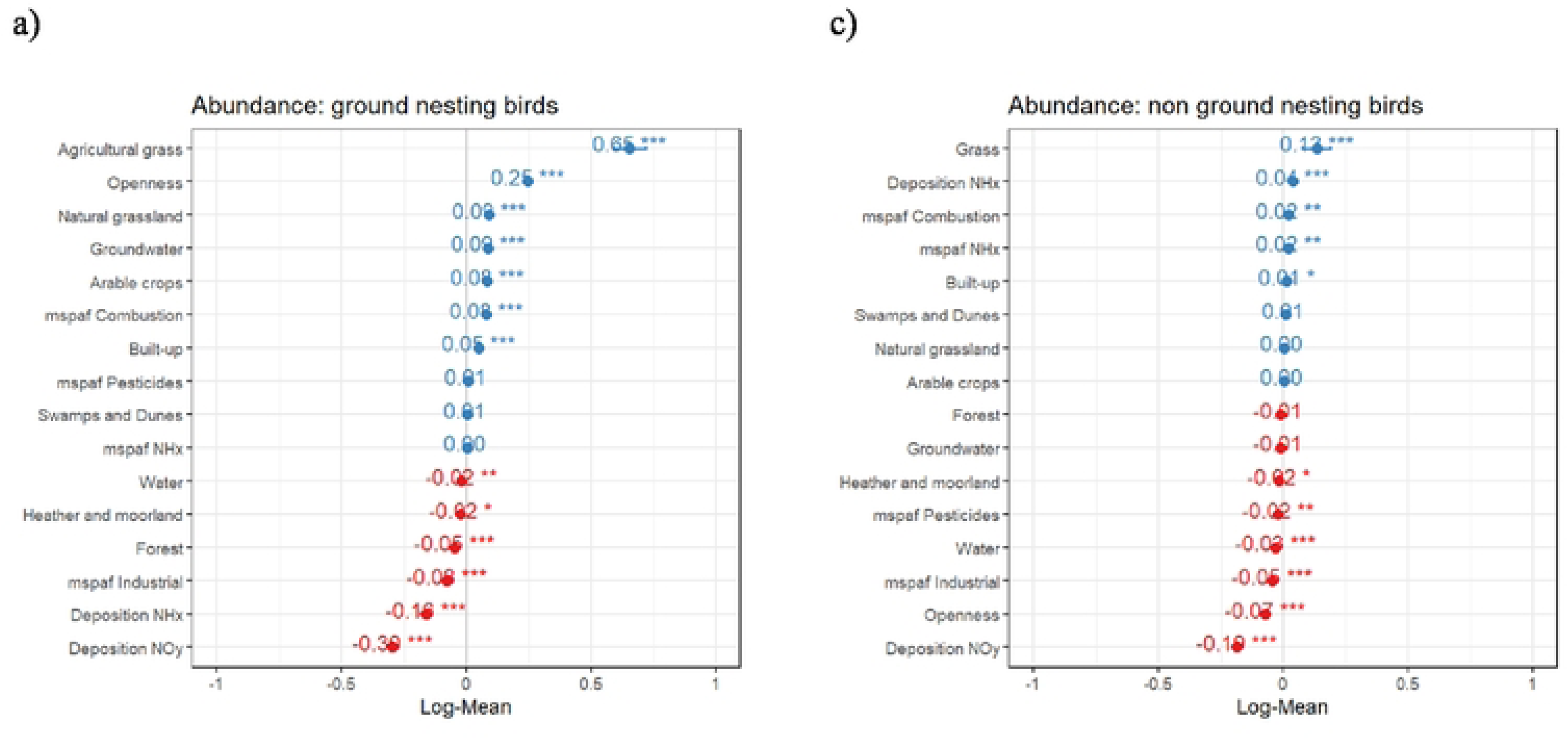
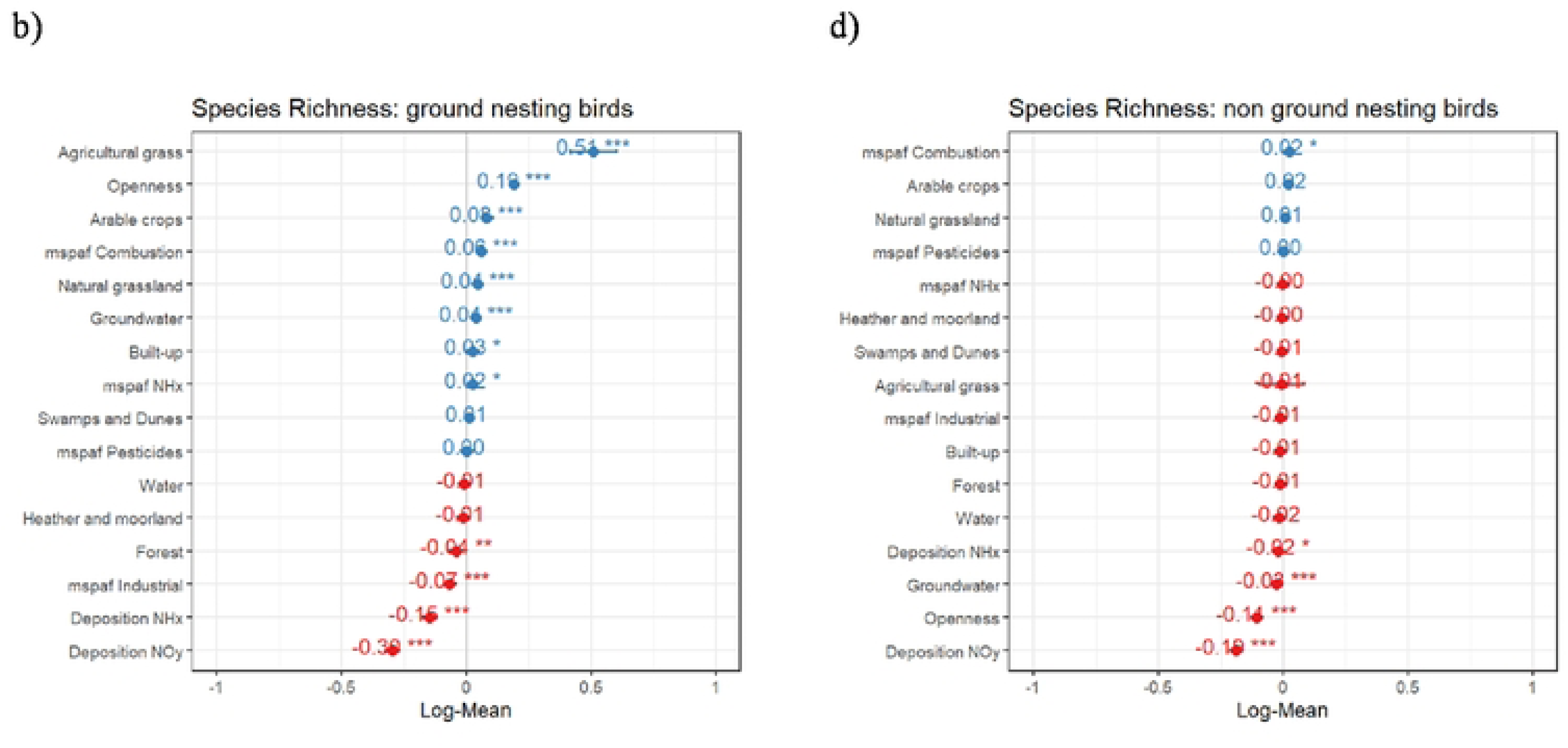
Relationship of farmland birds to landscape variable and pollutants in agricultural landscapes. Figure show results of model selection and averaging where the response variable is species richness or abundance of farmland birds (a,b) ground nesting birds; c,d) non-ground nesting birds) and land use, water and land pollutants (described in the text) are explanatory variables. Agricultural domains and survey type were included as random effects.

**Figure 3:**
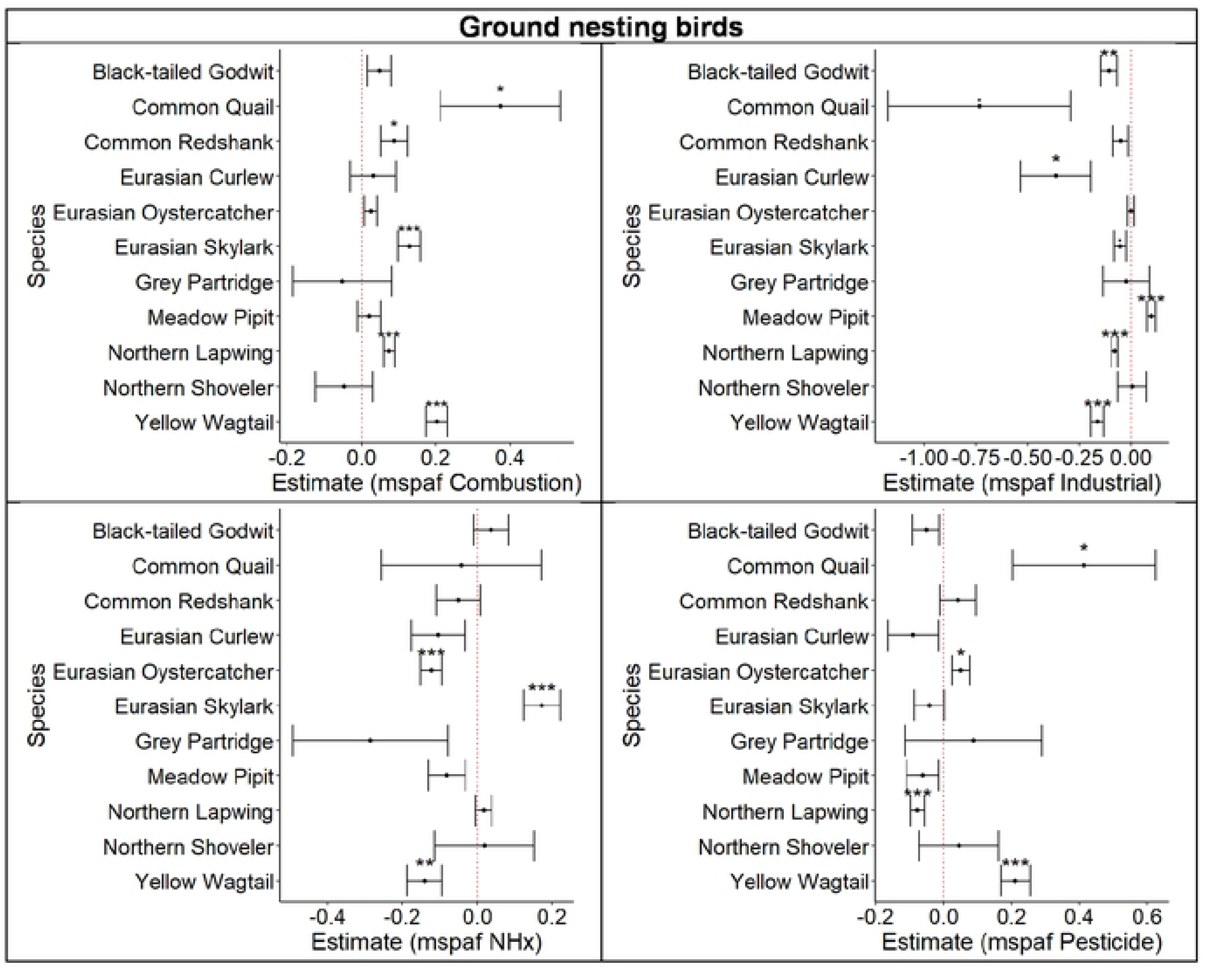
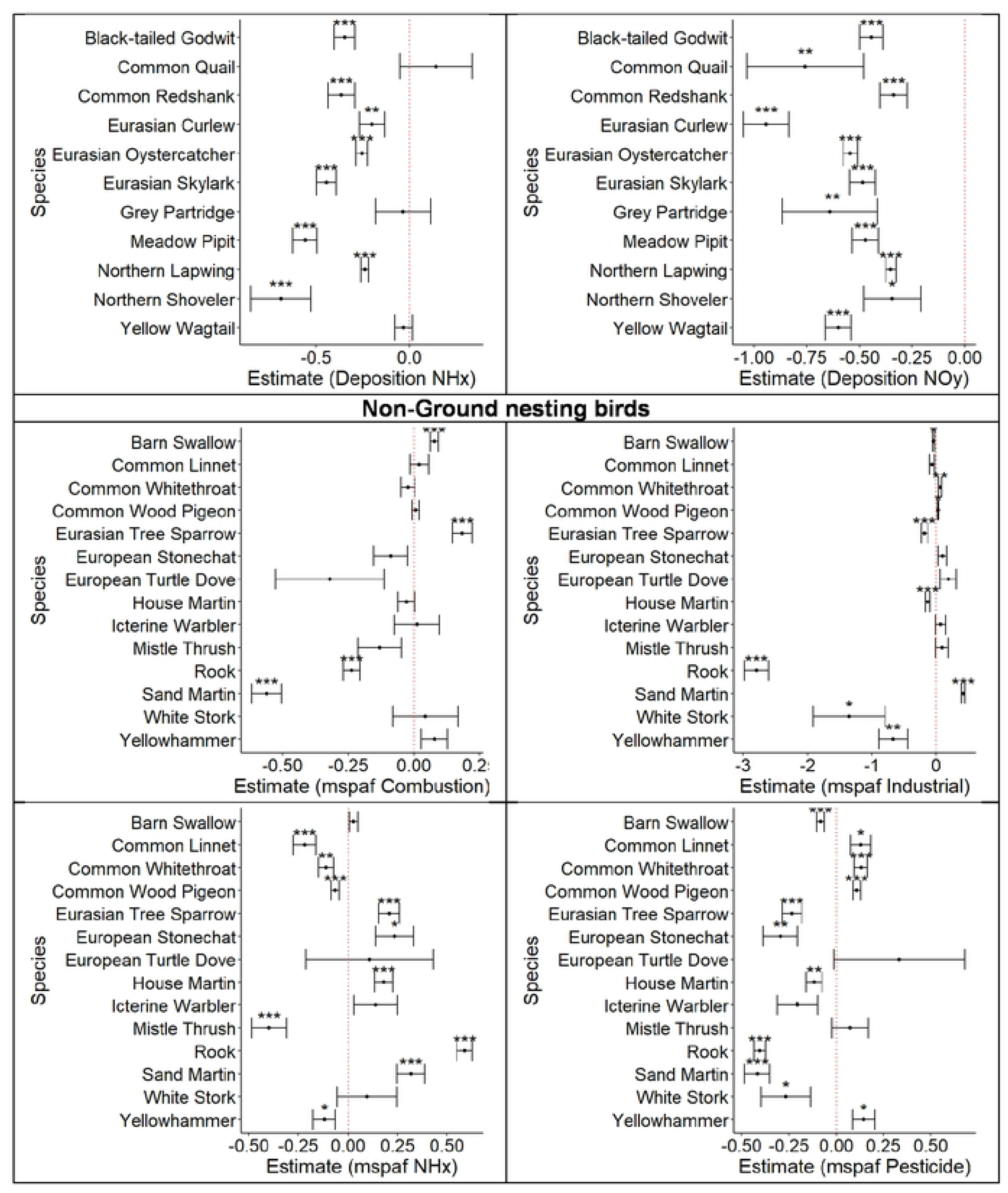
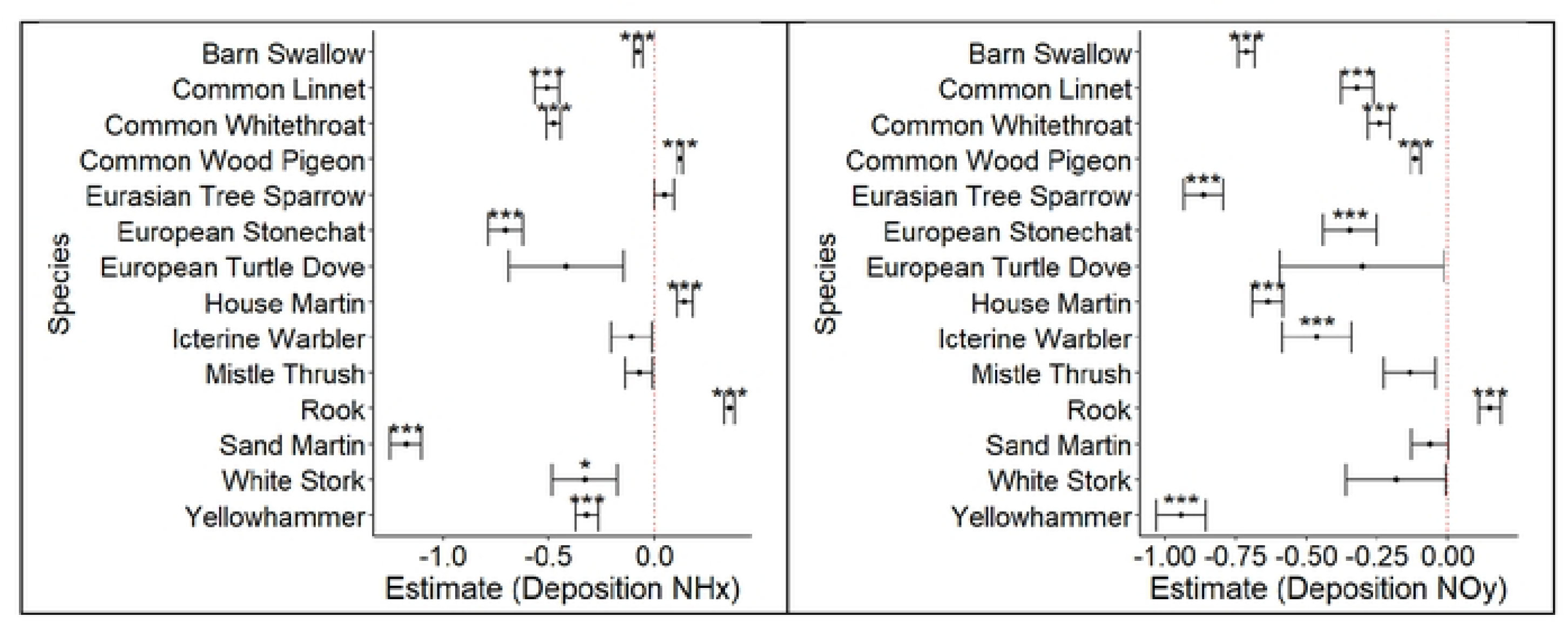
Response of individual species of farmland birds to surface water toxicity and atmospheric deposition. Results from individual species models were aggregated and ploted per pollutant. The significance of the results is represented by asterisks(*).

**Figure 4:**
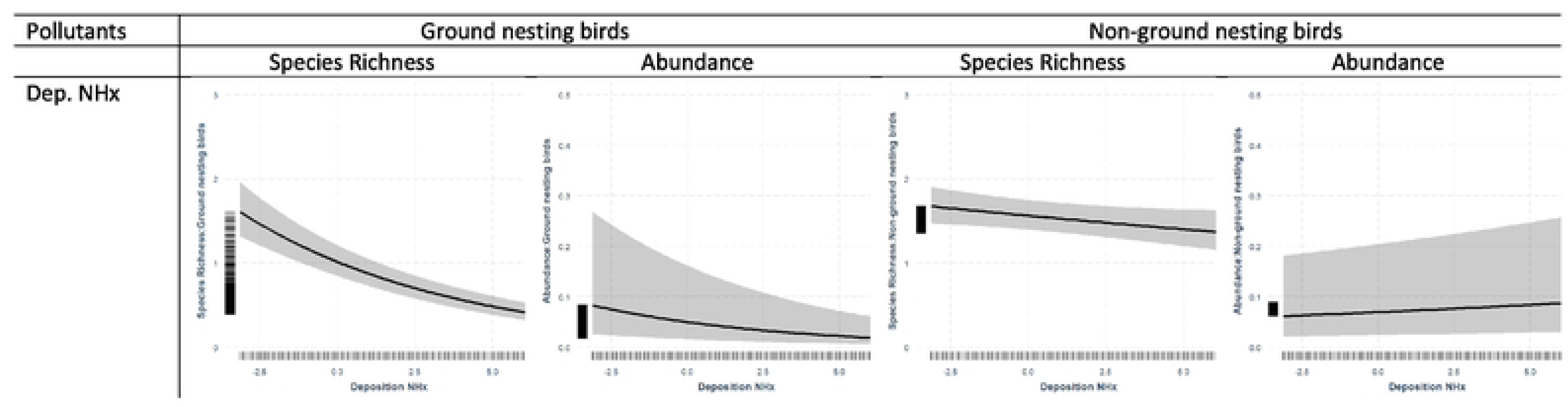
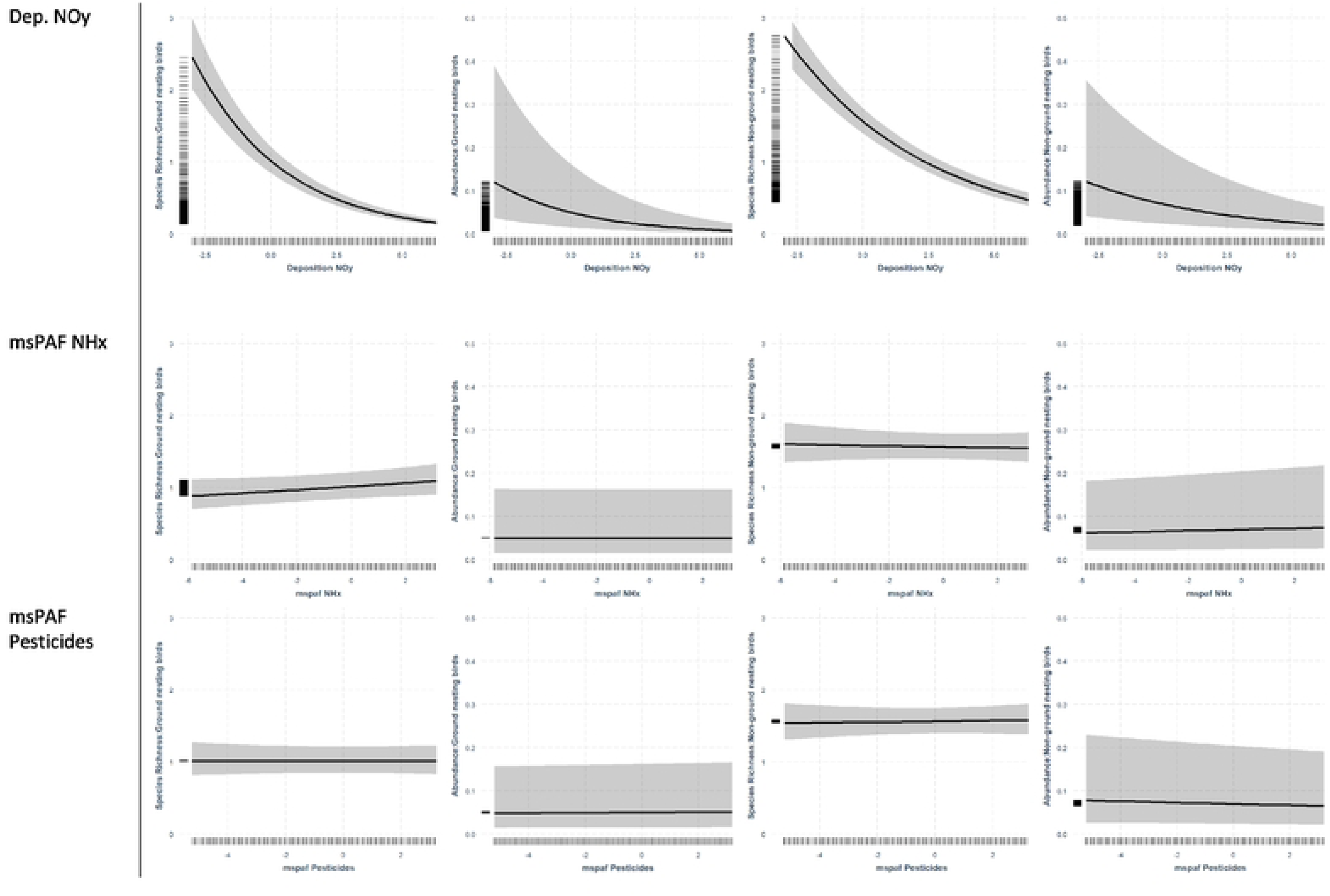
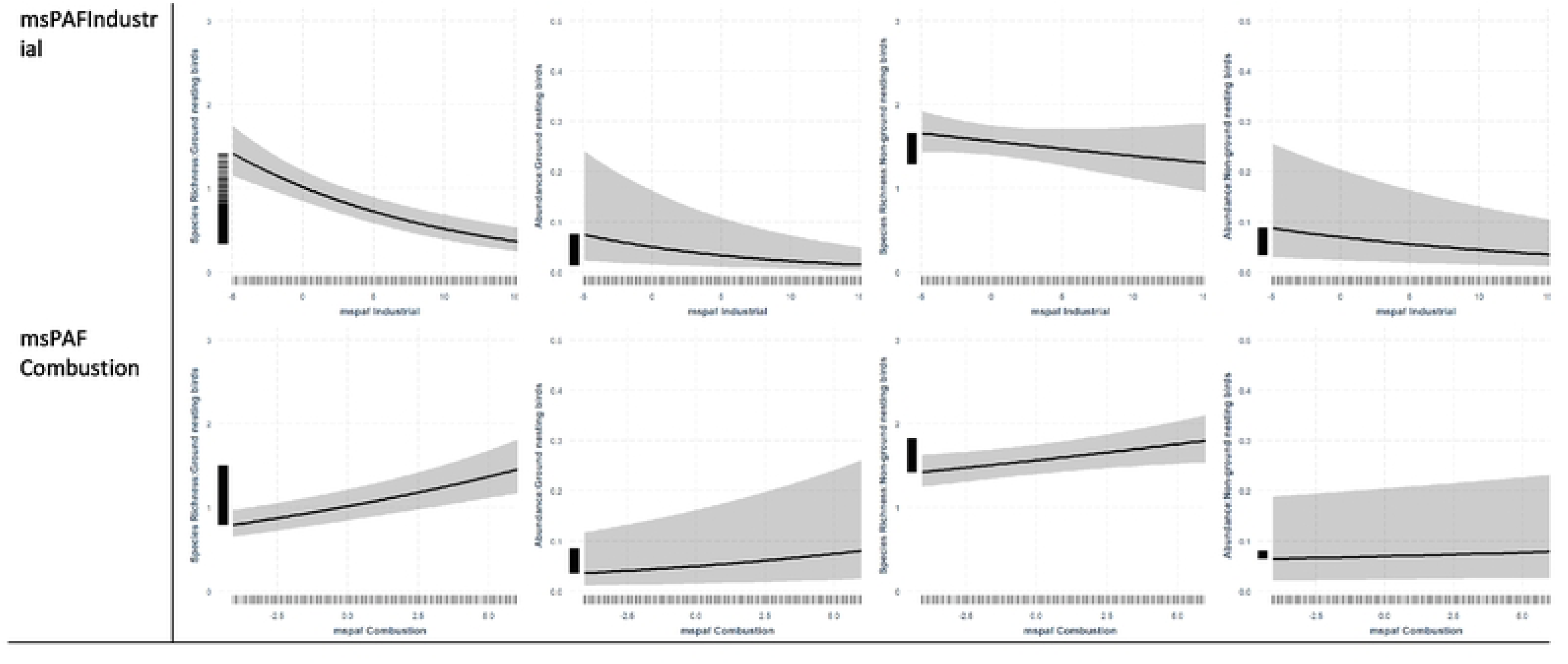
Modelled estimates of spe,:.ies richness and abundance of ground nesting and non-ground nesting farmland birds, as predicted by water pollutants and atmospheric deposition: Water pollutants used were mixture toxic pressure: msPAF NHx, msPAF pesticides, msPAF industrial, msPAF combustion and atmospheric deposition used were deposition NHx and deposition Noy. Variables were rescaled for modelling purposes (see text). Grey bands represent *95%* confidence intervals. A rug plot is also incorporated, displaying **individual data points along the axes for further context.**

The observed relationships between ground nesting farmland birds and the toxic pressure proxies were more varied and less consistent (Fig. 3,4). Ground nesting bird abundance and species richness were significantly negatively related to the mixture toxic pressure attributable to emissions of industrial chemicals, but significantly positively related to the mixture toxic pressure attributable to emissions from combustion processes. Ground nesting bird species richness was significantly positively related to the toxic pressure exerted by NHx. We found no support for relations with the mixture toxic pressure of pesticides in local surface waters, used as proxy for local exposure levels in the landscape.

#### 3.2.2 Non-ground-nesting bird species

Apart from clear negative relations with openness of the landscape, for abundance and species richness of non-ground-nesting birds, there was not much support for strong and consistent relations with land use variables (Fig.2b, SI8). This perhaps reflects the more variable composition of this group as compared to the ground-nesting species, that contains species that, in the Netherlands, preferentially nest in natural holes and nest boxes and buildings (tree sparrow), trees (rook) or shrubs (yellowhammer), alongside a notable diversity in migration strategies and diet.

In line with the results of the ground nesting birds, we found strong support for a negative relation between abundance and species richness of non-ground nesting birds and NOy deposition (Fig.3, 4). However, while the species richness of non-ground nesting birds was likewise negatively related, abundance was significantly positively related to deposition of NHx, although with a small effect size. Non-ground nesting bird abundance was furthermore significantly positively related to the mixture toxic pressure attributable to emissions from combustion processes and the toxic pressure exerted by NHx, while it was negatively related to the mixture toxic pressure attributable to use of pesticides and the mixture toxic pressure caused by emissions of industrial chemicals. Species richness of non-ground nesting birds was significantly positively related to the mixture toxic pressure of chemicals emitted from combustion processes.

The sensitivity analysis run on the subset of km grids that were within a 5 km radius of a measurement point for water toxicity showed results that were consistent with the findings of the overall analysis suggesting that the data interpolation did not affect the outcomes (SI9).

#### 3.2.3 Relations between pollutants and changes in species richness between 1998 and 2018

Between 1998 and 2018, overall species richness decreased by an average of two species per km^2^ (mean ± sd; ground nesting birds = −1.37 ± 1.82; an overall decrease of 46%; non-ground nesting birds = −3.17 ± 2.19; an overall decrease of 29%). Species richness of ground nesting birds declined faster in areas with a high proportion of water and grass and the decline in richness of non-ground nesting birds was more pronounced in non-open areas (Fig. 5). This was primarily the result of the fact that in 1998 farmland bird species richness in environmentally less suitable areas was already very low (Fig. 6) so that species richness could still decline only in areas which were more suitable for them. For the same reason we observed a positive relation between the change in species richness of ground-nesting birds and the deposition of NHx and NOy. This means that, for ground nesting birds in 1998, low deposition square km cells (<400 NOy mol ha-1 yr-1 or <500 NHx mol ha-1 yr-1; Fig. 6) still contained between 4 and 7 farmland bird species on average across domains. At the other extreme (>700 Noy mol ha-1 yr-1 or >2000 NHx mol ha-1 yr-1), 1 species per square km cell remained or species were already entirely absent in 1998. As opposed to ground nesting birds, non-ground nesting birds exhibited a weakly positively relationship with NOy deposition in 1998, and their decline was stronger with these higher NOy deposition loads resulting in an overall negative relation between NOy deposition and species richness (Fig. 5). With respect to the other pollutants, only mixture toxic pressure attributable to emissions from combustion processes was significantly positively related to the species richness trend of non-ground nesting birds.

**Figure 5:**
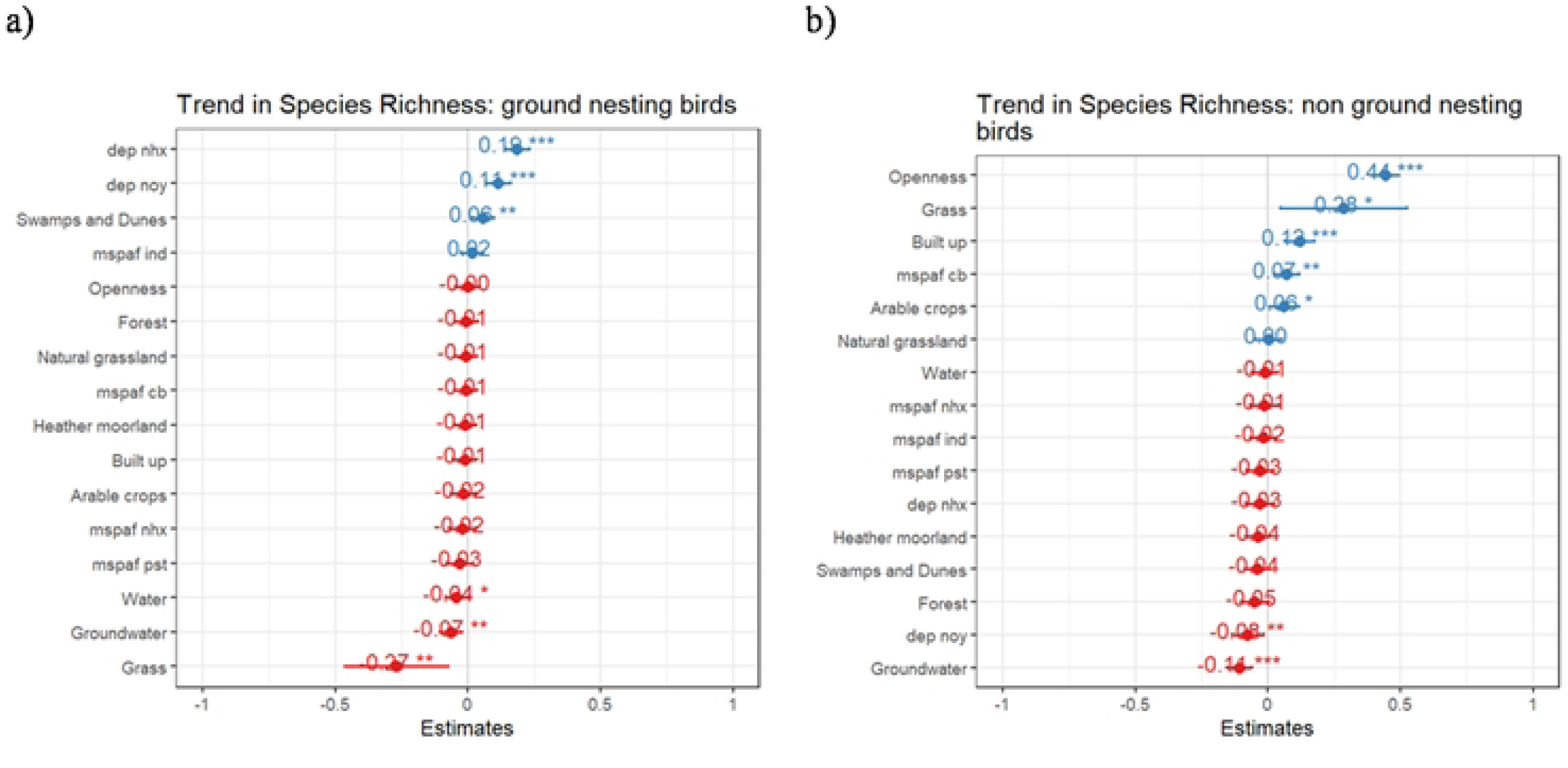
Relationship of farmland birds trends to landscape variable and pollutants in agricultural landscapes: The results of linear mixed effect models where the response variable is change in species richness from 1998 to 2008 of two groups of farmland birds ( a)ground nesting birds and b) non-ground nesting birds) and land use, water and land pollutants (described in the text) are explanatory variables. Agricultural domains and survey type were included as random effects.

**Figure 6:**
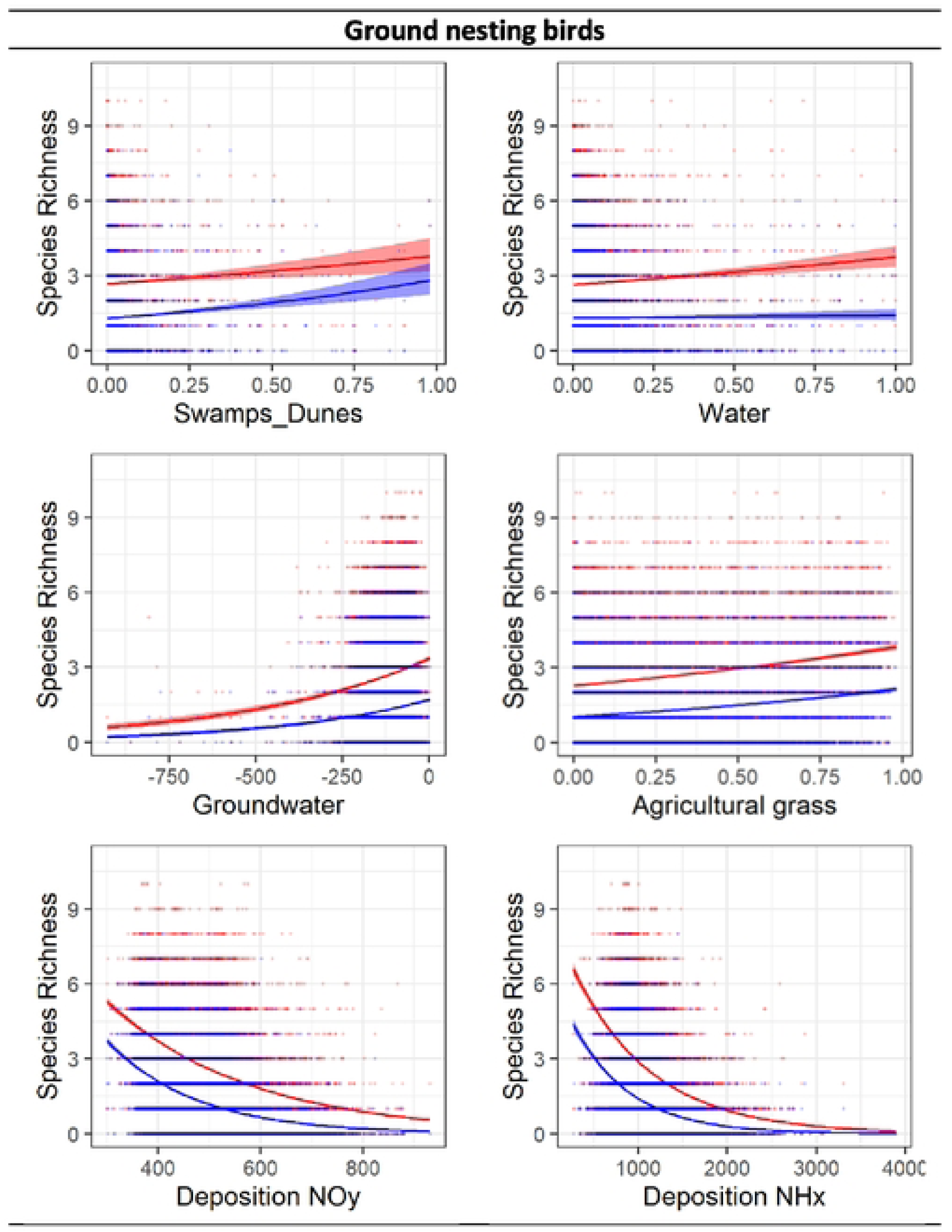
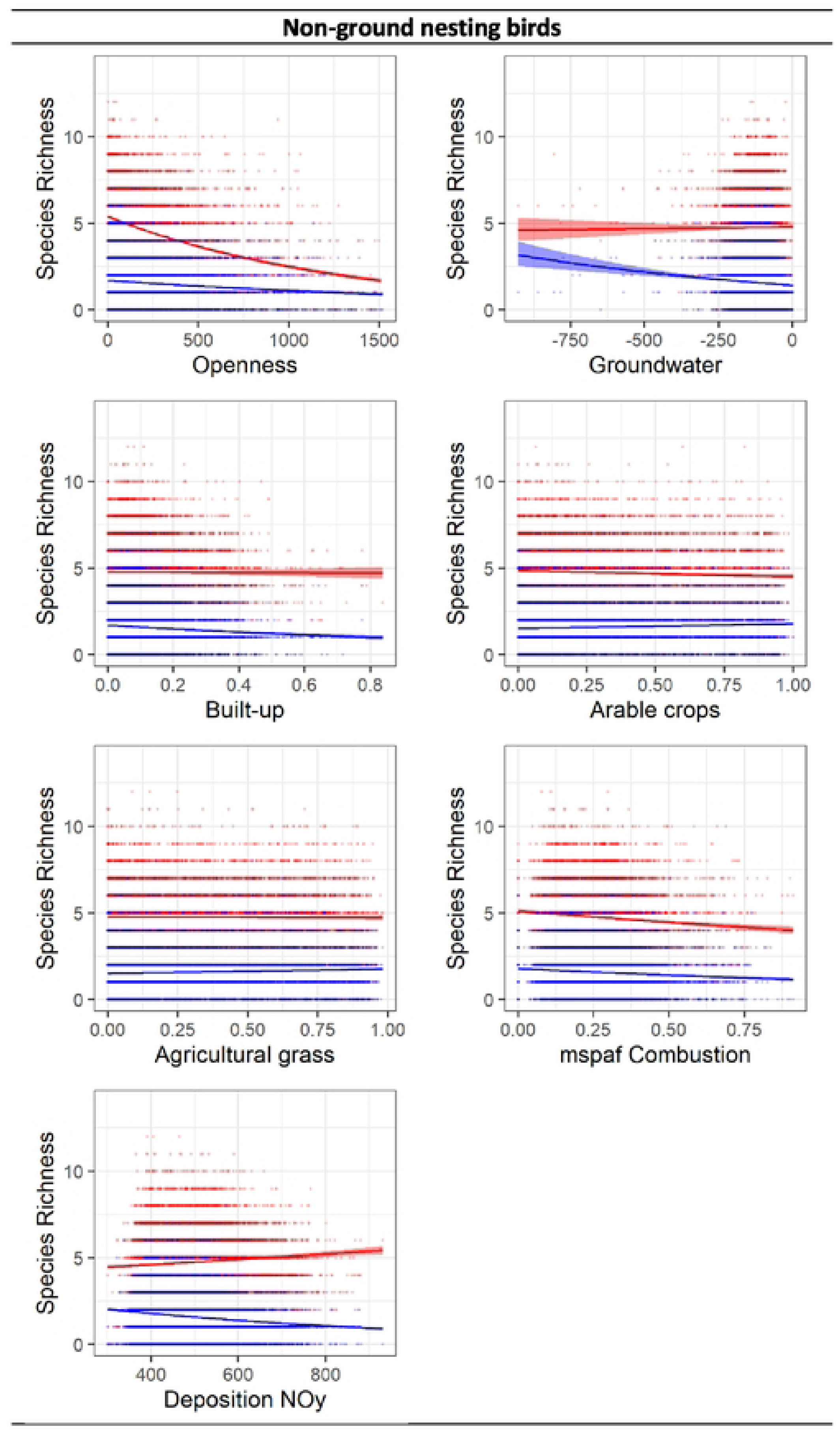
Modelled estimates of species richness and abundance trends of ground nesting and non-ground nesting farmland birds, as predicted by landscape variables and pollutants in agricultural landscapes. Figures show plots created using the ggsmooth function in R with a Poisson regression model, showing the relationship between species richness and abundance of fannland birds to all significant landscape variables and pollutants in agricultural landscapes from the results of the trend analysis shov.’Il in fig. 5. Data points and lines in red represent observations from the bird atlas from 1998, while those in blue signify data from 2018. The light red and light blue shaded areas represent 95% confidence intervals for the respective years. **A** rug plot is also incorporated, displaying individual data points along the axes for further context.

## 4 Discussion

There is major concern about the role of synthetic pollutants and acidifying eutrophicating substances in biodiversity decline, both alone and in combinations. Here, linking several datasets, unique in their nation-wide coverage and high spatial resolution, we investigated the relations between these pollutants and farmland birds, correcting for their distribution across the Netherlands and the landscape characteristics to which they adhere to. Most striking are the strongly negative relationships of farmland bird species richness and abundance with atmospheric deposition of NOy. Where at low deposition values around 5 species per 1×1 km2 grid cell can be encountered, at high deposition values farmland birds are essentially absent. The relationships between bird abundance and species richness and the metric to represent local toxic pressure levels in water were less pronounced and varied in direction and magnitude.

### 4.1 Farmland bird distribution in relation to landscape variables

We observed that the correlation between landscape variables and ground-nesting bird groups conformed to our expectations and aligned with established distribution patterns (Sovon, 2018). In the case of non-ground nesting birds, the relationships with landscape variables, such as openness, were as anticipated but displayed a weaker connection to other landscape variables. This discrepancy might be attributed to the diverse array of species within this group, vastly differing in feeding habits (from obligatory insectivores and seedeaters to facultative herbivores and carnivores) and migratory behavior. We also noted a slight difference in the relationship between landscape variables to abundance of non-ground nesting birds and the species richness of the same group. This difference may be attributed to a delayed response of species richness to changes in the landscape, in contrast to the more immediate response observed in abundance.

### 4.2 Farmland bird distribution in relation to nitrogen deposition

The models indicate a robust negative correlation between deposition of NOy and farmland bird populations. The distribution maps of the pollutants (Fig. 2a, b) showed a strong association between the presence of NOy and significant road networks and industrial activities in the Netherlands. This finding aligns with other studies indicating that in agricultural landscapes, the distribution of these urban pollutants, including NO_2_, is predominantly confined to areas in close proximity to major roads and airports. It remains plausible that NOy serves as an indicator of the presence of roads and industrial activities, both of which are recognized for their adverse effects on bird (nesting and breeding) densities (Cooke et al., 2020; Reijnen & Foppen, 2006).

We also found strong negative relations between deposition of NHx and ground nesting farmland birds and species richness of non-ground nesting birds. In the Netherlands, NHx deposition primarily originates from farming activities and livestock. In semi-natural and relatively nutrient-poor grasslands, NHx deposition exerts a substantial impact on vegetation by diminishing floral diversity and favoring the proliferation of generalist species (Dupre et al., 2009; Van Den Berg et al., 2010) that may negatively affect farmland biodiversity (plausible mechanisms discussed in Feest et al., 2014,Weiss, 1999). However, most agricultural grasslands in the Netherlands have been heavily fertilized since the 1970’s and it is unlikely that the relatively small added contribution of atmospheric nitrogen compounds could explain the observed relationships in Dutch farmland as found in our study.

A more likely explanation is that NHx deposition is an indicator of the intensity of farming in general, and is correlated with other agricultural practices such as fertilizer application amounts and grass cutting frequency, together with high sward densities (Kentie et al., 2015; Kleijn et al., 2010). Furthermore, the increased densities of grazing dairy cattle has led the conversion of meadows into monocultures of *Lolium perenne*, which, as a vegetation type, is highly unfavorable for meadow birds

### 4.3 Farmland bird distribution in relation to local toxicity estimates

The relationships between farmland bird diversity and abundance with local estimates of toxic pressure, approximated via local surface water toxicity metrics, were weaker and more variable than with atmospheric nitrogen deposition. It is conceivable that this discrepancy may be partially attributed to differences in resolution between the data from the deposition maps and the resolution attainable through kriging maps generated from the water monitoring points. Such differences in resolution could potentially influence the precision of the true local exposure levels, and thus the attribution of effect sizes to this pressure type as observed in the model results. It may also explain the stronger effects of atmospheric NHx relative to those of mspaf NHx (Fig. 4, SI8).

Although the influence of toxic pressure on farmland birds could only be characterized as relatively mild with the available data, there were some interesting insights. First, we found a consistent positive relationship between abundance and species richness of both groups of farmland birds and mixture toxic pressure attributable to emissions from combustion processes. While this may be associated with landscape features and other indirect effects, the exact cause for this relationship remains obscure.

Second, although our comprehensive assessment of mixture toxic pressure levels attributable to the use of pesticides (mixture toxic pressure attributable to pesticides) did not yield consistent findings, other studies in the Netherlands have revealed substantial negative effects on the trends of farmland birds linked to elevated surface water concentrations of specific pollutants like imidacloprid (Hallmann et al., 2014). One plausible reason for this could be that unlike the study by Hallman et al. (2014) where only insectivores were considered, our study included species with a more diverse diet type. Species with a more herbivorous diet may be less sensitive as pesticides accumulate in the food chain.

Interestingly we found a consistent negative impact of local toxic pressure levels, due to exposures caused by emissions of industrial pollutants (msPAF industrial) and farmland birds belonging to both groups. Multiple research investigations have determined that industrial contaminants, including polycyclic aromatic hydrocarbons (PAHs) and polychlorinated biphenyl (PCB), have the potential to impact both the production and quality of eggs, accumulated through the food items of female parents. This influence can result in increased instances of hatching failure, characterized by elevated embryonic mortality and developmental abnormalities (Albers, 2006; Fernie et al., 2000; Koeman et al., 1973). The sources of these industrial pollutants in the hydrological systems in agricultural lands, where they were measured, include atmospheric deposition, organic amendments like manure, and runoff from impervious surfaces. These can contaminate soil and crops, becoming part of the farmland bird food chain.

### 4.4 Changes in distribution patterns in recent years

Although there has been no profound transformation in the main landscape types in the Netherlands between 1998 and 2018, our results (Fig. 6) show a substantial overall decline in bird species richness across the Netherlands. The trend analysis revealed that most of the relationships between bird distribution and nitrogen deposition were already apparent 20 years ago (also see Lenon et al., 2019). Recent declines occurred for the most part in landscapes that harbored a fairly large number of species, such as in low nitrogen deposition grid cells. Similarly, ground nesting birds that typically favor agricultural grasslands showed a more pronounced decrease in species richness in 2018 than in 1998. This corresponds with the observations that in livestock grassland-based systems, there is a negative trend in associated biodiversity, due to more frequent and earlier mowing of grasslands, high nutrient inputs, and lowered groundwater table, collectively contributing to lower diversity (more details in Brink, 2015). It is conceivable that pollutants serve as indicators of broader landscape changes, making it challenging to untangle the associated factors and correlations among variables that may have not been considered in this analysis.

## 5 Conclusion

Aerial pollutants such as NH_3_, NO_2_, and SO_2_ can compromise bird health, behavior and reproductive success (Fernie et al., 2016; Richard et al., 2021; Salmón et al., 2018; Sanderfoot & Holloway, 2017). The consistent negative relationships that we find between nitrogen (N) deposition and farmland bird diversity and abundance are consistent with these direct negative effects. However, it is likely that these relationships are a proxy for the intensity of farming and human disturbance in general. Moreover, our study reveals diverse and mixed associations between farmland birds and local toxic pressure variation as established via measured exposures in surface waters, with consistent relationships for the mixture toxic pressure caused by exposure to industrial chemicals (negative) and or to products of combustion (positive). The correlative evidence we provide, based on a nationwide unique integration of detailed datasets on farmland bird distribution, nitrogen deposition and water pollutants, calls for well-designed field studies to enhance the causal understanding of the patterns observed, as well as the design of appropriate measures in land use and pollution reduction.

## ACKNOWLEDGEMENT

This study was funded by the project “Transition to a sustainable food system” (with project number NWA.1235.18.201) which is financed by the Dutch Research Council (NWO). We also acknowledge NWO grant ALWPP.2019.007. We are grateful to the many thousands of volunteer fieldworkers for their invaluable effort in collecting data for the breeding bird atlas by counting birds throughout the Netherlands. LP and JS were funded by RIVM’s strategic research program, run under the auspices of RIVM’s Scientific Advisory Board, and STOWA (Dutch Foundation for Applied Water Research).

## AUTHOR CONTRIBUTIONS

All authors were part of the conceptualization and planning of the study design. Henk Sierdsema provided curated bird data for this study. Jaap Slootweg and Leo Posthuma supplied curated data and code for the characterization of toxic pressure of surface water. Maitreyi Sur was responsible for the collection of all other data and data analysis. Maitreyi Sur, David Kleijn, Hans de Kroon, and Merel Soons led the writing of the manuscript. All authors contributed critically to the drafts and gave final approval for publication.

## DATA AVAILABILITY

Data on farmland birds can be obtained from Sovon upon request, while information on surface water toxicity is available from RIVM upon request. All other data were sourced from publicly available resources described in the main text.

## SUPPORTING INFORMATION

**SI1:** Grids used for monitoring of farmland bird species in the Netherlands

**SI2**: Extra details of explanatory variables used in the analysis

**SI3**: Map showing distribution of 21 domains within the Netherlands.

**SI4**: Description of consolidated 9 classes of land use

**SI5**: Maps showing average openness of a landscape and average spring groundwater level

**SI6**: Table showing VIF values for four different models with model structures

**SI7**: Figure showing correlation matrix of explanatory variables

**SI8**: Model averaged results

**SI9**: Results from the sensitivity analysis

## REFERENCES

Aebischer, N. J., & Ewald, J. A. (2004). Managing the UK Grey Partridge Perdix perdix recovery: population change, reproduction, habitat and shooting. Ibis, 146, 181–191.

Albers, P. H. (2006). Birds and polycyclic aromatic hydrocarbons. Avian and Poultry Biology Reviews, 17(4), 125–140.

Altenburg, J., van Diek, H., Foppen, R., Kampichler, C., Sierdsema, H., Troost, G., van Winden, E., & van Turnhout, C. (2017). Fieldwork completed for the fourth Dutch bird atlas, a bonanza of counts and estimates to be utilised. Vogelwelt, 137, 23–28.

Barton, M. G., Henderson, I., Border, J. A., & Siriwardena, G. (2023). A review of the impacts of air pollution on terrestrial birds. Science of the Total Environment, 873, 162136.

Bibby, C. J., Burgess, N. D., & Hill, D. A. (2012). Bird Census Techniques. Elsevier Science.

Bobbink, R. (1991). Effects of nutrient enrichment in Dutch chalk grassland. Journal of applied ecology, 28–41.

Brickle, N. W., & Harper, D. G. (2002). Agricultural intensification and the timing of breeding of Corn Buntings Miliaria calandra. Bird study, 49(3), 219–228.

Brickle, N. W., Harper, D. G., Aebischer, N. J., & Cockayne, S. H. (2000). Effects of agricultural intensification on the breeding success of corn buntings Miliaria calandra. Journal of applied ecology, 37(5), 742–755.

Brink, M. (2015). Country Report for The State of the World’s Biodiversity for Food and Agriculture–The Netherlands: CGN Report 34.

Campbell, B. M., Beare, D. J., Bennett, E. M., Hall-Spencer, J. M., Ingram, J. S., Jaramillo, F., Ortiz, R., Ramankutty, N., Sayer, J. A., & Shindell, D. (2017). Agriculture production as a major driver of the Earth system exceeding planetary boundaries. Ecology and society, 22(4).

Cellier, P., Durand, P., Hutchings, N., Dragosits, U., Theobald, M., Drouet, J.-L., Oenema, O., Bleeker, A., Breuer, L., & Dalgaard, T. (2011). Nitrogen flows and fate in rural landscapes.

Cooke, S. C., Balmford, A., Donald, P. F., Newson, S. E., & Johnston, A. (2020). Roads as a contributor to landscape-scale variation in bird communities. Nature communications, 11(1), 3125.

Czyżewski, B., Trojanek, R., Dzikuć, M., & Czyżewski, A. (2020). Cost-effectiveness of the common agricultural policy and environmental policy in country districts: Spatial spillovers of pollution, bio-uniformity and green schemes in Poland. Science of the Total Environment, 726, 138254.

Dise, N. (2011). The European Nitrogen Assessment: Sources, effects and policy perspectives, chap. Nitrogen as a threat to European terrestrial biodiversity 700. In: Cambridge University Press.

Donald, P. F., Green, R., & Heath, M. (2001). Agricultural intensification and the collapse of Europe’s farmland bird populations. Proceedings of the Royal Society of London. Series B: Biological Sciences, 268(1462), 25–29.

Duprè, C., Stevens, C. J., Ranke, T., Bleeker, A., Peppler-Lisbach, C., Gowing, D. J., Dise, N. B., Dorland, E., Bobbink, R., & Diekmann, M. (2010). Changes in species richness and composition in European acidic grasslands over the past 70 years: the contribution of cumulative atmospheric nitrogen deposition. Global Change Biology, 16(1), 344–357.

Emmerson, M., Morales, M. B., Oñate, J. J., Batary, P., Berendse, F., Liira, J., Aavik, T., Guerrero, I., Bommarco, R., & Eggers, S. (2016). How agricultural intensification affects biodiversity and ecosystem services. In Advances in ecological research (Vol. 55, pp. 43–97). Elsevier.

Engel, J., Huth, A., & Frank, K. (2012). Bioenergy production and Skylark (Alauda arvensis) population abundance – a modelling approach for the analysis of land-use change impacts and conservation options. GCB Bioenergy, 4(6), 713–727.

Environmental Data Compendium. Trend fauna - all species monitored - Living Planet Index Netherlands, 1990-2021 (indicator 1569, version 08, 29 March 2023).

Feest, A., van Swaay, C., & van Hinsberg, A. (2014). Nitrogen deposition and the reduction of butterfly biodiversity quality in the Netherlands. Ecological Indicators, 39, 115–119.

Fernie, K. J., Bortolotti, G. R., Smits, J. E., Wilson, J., Drouillard, K. G., & Bird, D. M. (2000). Changes in egg composition of American kestrels exposed to dietary polychlorinated biphenyls. *Journal of Toxicology and Environmental Health*, Part A, 60(4), 291–303.

Fernie, K. J., Cruz-Martinez, L., Peters, L., Palace, V., & Smits, J. E. (2016). Inhaling benzene, toluene, nitrogen dioxide, and sulfur dioxide, disrupts thyroid function in captive American kestrels (Falco sparverius). Environmental Science & Technology, 50(20), 11311–11318.

Gregory, R. D., & van Strien, A. (2010). Wild bird indicators: using composite population trends of birds as measures of environmental health. Ornithological Science, 9(1), 3–22.

Hallmann, C. A., Foppen, R. P., Van Turnhout, C. A., De Kroon, H., & Jongejans, E. (2014). Declines in insectivorous birds are associated with high neonicotinoid concentrations. Nature, 511(7509), 341–343.

Humann-Guilleminot, S., Clément, S., Desprat, J., Binkowski, Ł. J., Glauser, G., & Helfenstein, F. (2019). A large-scale survey of house sparrows feathers reveals ubiquitous presence of neonicotinoids in farmlands. Science of the Total Environment, 660, 1091–1097.

IPBES. (2019). Global assessment report on biodiversity and ecosystem services of the Intergovernmental Science-Policy Platform on Biodiversity and Ecosystem Services.

Kentie, R., Both, C., Hooijmeijer, J. C., & Piersma, T. (2015). Management of modern agricultural landscapes increases nest predation rates in Black-tailed Godwits L imosa limosa. Ibis, 157(3), 614–625.

Kleijn, D., Schekkerman, H., Dimmers, W. J., Van Kats, R. J., Melman, D., & Teunissen, W. A. (2010). Adverse effects of agricultural intensification and climate change on breeding habitat quality of Black-tailed Godwits Limosa l. limosa in the Netherlands. Ibis, 152(3), 475–486.

Kleijn, D., & van Zuijlen, G. J. (2004). The conservation effects of meadow bird agreements on farmland in Zeeland, The Netherlands, in the period 1989–1995. Biological Conservation, 117(4), 443–451.

Koeman, J., Van Velzen-Blad, H., De Vries, R., & Vos, J. (1973). Effects of PCB and DDE in cormorants and evaluation of PCB residues from an experimental study. J Reprod Fert Suppl, 19, 353–364.

Köhler, H.-R., & Triebskorn, R. (2013). Wildlife ecotoxicology of pesticides: can we track effects to the population level and beyond? science, 341(6147), 759–765.

Kwak, R., & Louwe Kooijmans, J. (2021). Nederlandse vogels in hun domein. (Vogelbescherming Nederland). KNNV Uitgeverij

Lennon, R.J., Isaac, N.J., Shore, R.F., Peach, W.J., Dunn, J.C., Pereira, M.G., Arnold, K.E., Garthwaite, D. & Brown, C.D. (2019). Using long-term datasets to assess the impacts of dietary exposure to neonicotinoids on farmland bird populations in England. PloS one, 14(*10*), p.e0223093.

Li, Y., Miao, R., & Khanna, M. (2020). Neonicotinoids and decline in bird biodiversity in the United States. Nature Sustainability, 3(12), 1027–1035.

Maskell, L. C., Smart, S. M., Bullock, J. M., Thompson, K., & Stevens, C. J. (2010). Nitrogen deposition causes widespread loss of species richness in British habitats. Global Change Biology, 16(2), 671–679.

Mauser, W., Klepper, G., Zabel, F., Delzeit, R., Hank, T., Putzenlechner, B., & Calzadilla, A. (2015). Global biomass production potentials exceed expected future demand without the need for cropland expansion. Nature communications, 6(1), 8946.

Moreau, J., Rabdeau, J., Badenhausser, I., Giraudeau, M., Sepp, T., Crépin, M., Gaffard, A., Bretagnolle, V., & Monceau, K. (2022). Pesticide impacts on avian species with special reference to farmland birds: a review. Environmental Monitoring and Assessment, 194(11), 790.

Morris, A. J., Wilson, J. D., Whittingham, M. J., & Bradbury, R. B. (2005). Indirect effects of pesticides on breeding yellowhammer (Emberiza citrinella). Agriculture, ecosystems & environment, 106(1), 1–16.

Newbold, T., Hudson, L. N., Hill, S. L., Contu, S., Lysenko, I., Senior, R. A., Börger, L., Bennett, D. J., Choimes, A., & Collen, B. (2015). Global effects of land use on local terrestrial biodiversity. Nature, 520(7545), 45–50.

Newton, I. (2004). The recent declines of farmland bird populations in Britain: an appraisal of causal factors and conservation actions. Ibis, 146(4), 579–600. 10.1111/j.1474-919X.2004.00375.x

Nijssen, M., WallisDeVries, M., & Siepel, H. (2017). Pathways for the effects of increased nitrogen deposition on fauna. Biological Conservation, 212, 423–431.

OECD. (2023). Agricultural land (indicator). doi: 10.1787/9d1ffd68-en *(Accessed on* 05 July 2023)

Reijnen, R., & Foppen, R. (2006). Impact of road traffic on breeding bird populations. In The ecology of transportation: managing mobility for the environment (pp. 255-274). Springer.

Richard, F.-J., Gigauri, M., Bellini, G., Rojas, O., & Runde, A. (2021). Warning on nine pollutants and their effects on avian communities. Global Ecology and Conservation, 32, e01898.

Rigal, S., Dakos, V., Alonso, H., Auniņš, A., Benkő, Z., Brotons, L., Chodkiewicz, T., Chylarecki, P., de Carli, E., del Moral, J. C., Domşa, C., Escandell, V., Fontaine, B., Foppen, R., Gregory, R., Harris, S., Herrando, S., Husby, M., Ieronymidou, C., . . . Devictor, V. (2023). Farmland practices are driving bird population decline across Europe. Proceedings of the National Academy of Sciences, 120(21), e2216573120.

Salmón, P., Stroh, E., Herrera-Dueñas, A., von Post, M., & Isaksson, C. (2018). Oxidative stress in birds along a NOx and urbanisation gradient: an interspecific approach. Science of the Total Environment, 622, 635–643.

Sanderfoot, O. V., & Holloway, T. (2017). Air pollution impacts on avian species via inhalation exposure and associated outcomes. Environmental Research Letters, 12(8), 083002.

Sovon. (2018). Vogelatlas van Nederland.

Stevens, C., Bell, J., Brimblecombe, P., Clark, C., Dise, N., Fowler, D., Lovett, G., & Wolseley, P. (2020). The impact of air pollution on terrestrial managed and natural vegetation. Philosophical Transactions of the Royal Society A, 378(2183), 20190317.

Sutton, M. A., Howard, C. M., Erisman, J. W., Billen, G., Bleeker, A., Grennfelt, P., Van Grinsven, H., & Grizzetti, B. (2011). The European nitrogen assessment: sources, effects and policy perspectives. Cambridge University Press.

Tang, F. H., Lenzen, M., McBratney, A., & Maggi, F. (2021). Risk of pesticide pollution at the global scale. Nature Geoscience, 14(4), 206–210.

Tilman, D., Balzer, C., Hill, J., & Befort, B. L. (2011). Global food demand and the sustainable intensification of agriculture. Proceedings of the National Academy of Sciences, 108(50), 20260–20264.

Van den Berg, L. J., Vergeer, P., Rich, T. C., Smart, S. M., Guest, D., & Ashmore, M. R. (2011). Direct and indirect effects of nitrogen deposition on species composition change in calcareous grasslands. Global Change Biology, 17(5), 1871–1883.

Vorisek, P., Klvanova, A., Wotton, S. & Gregory, R.D. (2008). A best practice guide for wild bird monitoring schemes. Praga: CSO/RSPB.

Weiss, S. B. (1999). Cars, cows, and checkerspot butterflies: nitrogen deposition and management of nutrient-poor grasslands for a threatened species. Conservation Biology, 13(6), 1476–1486.

